# Modeling of the Network Mediated by IL-36 Involved in Psoriasis

**DOI:** 10.1101/2023.07.03.547538

**Authors:** Sneha Pandey, Syona Tiwari, Sulagna Basu, Rajiv Kumar Mishra, Rakesh Pandey

## Abstract

Pathogenesis of inflammatory, chronic and common skin disease Psoriasis involves immune cells, skin cells (keratinocytes) and cytokines secreted by them. Hyperproliferation and abnormal differentiation of keratinocytes are believed to be a hallmark of it. Roles of several cytokines such as TNF*α*, IL-15, IL-17 and IL-23 in Psoriasis have been explored through mathematical/computational models as well as experimentally. However, the role of pro-inflammatory cytokine IL-36 is still elusive, especially in the case of General Pustular Psoriasis, a prevalent type of Psoriasis. To explore the role of that here, we construct a network embodying indirect cell-cell interactions of a few immune and skin cells mediated by IL-36 based on the existing knowledge. Further, we develop a mathematical model for the network and study the steady-state behaviour of that. Our results demonstrate that an increase in the level of IL-36 could lead to the hyper-proliferation of keratinocytes and, thus, Psoriasis. In addition, the analysis suggests that the plaque formation and progression of Psoriasis could occur via a gradual or switch-like increase in the population of keratinocytes. The switch-like increase would be due to the bistable behaviour of the network and could be used as a novel treatment strategy, as proposed and demonstrated earlier.

## 1 Introduction

Psoriasis is one of the common skin diseases, and approximately 2-3 % of the world population [1] is affected by it. It is inflammatory and chronic in nature, distinguished by the hyper-proliferation of keratinocytes (skin cells) and infiltration of immune cells in the Psoriatic lesions. The lesion could be scaly or in the form of pustules characterising different types of psoriasis. The involvement of Immune cells such as T-lymphocytes, dendritic, macrophages, mast and neutrophil cells in the pathophysiology of Psoriasis has been well-established[2]. The interaction among these immune and skin cells is believed to be mediated by cytokines secreted by them. Recently, roles of cytokines TNF*α*, IL-15, and IL-17/IL-23 axis have been demonstrated by constructing networks of indirect cell-cell interactions among immune cells and keratinocytes mediated by them [3][4]. Based on the steady-state behaviour of the mathematical models of these networks, authors have demonstrated how an increase in the level of modelled cytokines could lead to the progression of Psoriasis. Cytokines involved in Psoriasis vulgaris have been extensively explored compared to the other types of Psoriasis, such as Generalised Pustular Psoriasis (GPP) in which inflammatory cytokine IL-36 is believed to have a key role [5]. However, the underlying mechanism of its role in the pathogenesis of Psoriasis is still elusive. To address that, we have constructed a network for the indirect cell-cell interactions among immune and keratinocyte cells mediated by the cytokine IL-36. Our network is built on the published observations about the interactions of epidermal keratinocytes, T cells, macrophages, and dendritic cells. Modulation of cells through IL-36 occurs upon binding of IL-36 on its receptors present on the surface of modelled cells in the network. Keratinocytes (K) are a predominant source of IL-36 cytokines, and in an autocrine manner, it enhances the expression of IL-36 [6]. In case of a skin injury in presence of Psoriasis stimulates the release of IL-36 by neighbouring keratinocytes in a paracrine manner by releasing cathelicidin LL-37 from dead keratinocytes[7, 8]. Due to that, T cells and dendritic cells are recruited at the injury site [9]. The IL-36 receptors are predominantly present on naive *CD*4^+^ T cells, and one of the isoforms of that (IL36*β*) is constitutively expressed in them [10]. Many studies have reported that these *CD*4^+^ T cells, present in the psoriatic lesion, hyper proliferate in response to an IL-36 stimulus [11]. It has been observed that the bone marrow-derived dendritic cells show Th1 responses as a result of IL-36 induction [10]. The DCs activated by IL-36 secrete IL-12, and a synergistic effect of both the cytokines results in the polarization of naive *CD*4^+^*T* cells towards Th1 [10]. Also, IL-36 promotes the Th17 and Th22 lymphocyte polarisation through a similar DC-mediated mechanism [9, 12]. It is also observed that the IL-36-mediated T cell activation by dendritic cells leads to the hyperproliferation of keratinocytes[13].

Macrophages are also one of the sources of IL-36 cytokine in a Psoriatic lesion[14]. Macrophages treated with IL36*γ* stimulate the release of TNF*α* and IL-23 that have multiple roles such as in differentiation and proliferation of T cells[15], in the induction of cytokines by KCs (illustrated by [16]) and in the maturation of DCs. Human dermal macrophages have a high concentration of IL-36R on their surface, which is converted into a pro-inflammatory form (M1 phenotype) upon the action of IL-36 cytokines[14]. Cytokine-stimulated macrophages increase the adherence of monocytes on endothelial cells and cause the polarization of lymphocytes (here, T cell) due to the upregulation of the IL17/IL22 axis [15].

Here, we have developed a mathematical model for the constructed network mediated by IL-36 to explore the steady-state behaviour of the network. Our results demonstrate that an increase in the level of IL-36 could lead to the hyper-proliferation of keratinocytes which is considered as a hallmark of Psoriasis. We also observed that the route of the plaque formation and or progression of Psoriasis could be via a gradual or switch-like increase in the population of keratinocytes. This switch-like behaviour could be used as a new treatment strategy, as argued in a few recent studies [4, 17].

## 2 Materials and Methods

Based on the available published knowledge about the role of cytokine IL-36 in the pathogenesis of Psoriasis and General Pustular Psoriasis, a schematic diagram is prepared to depict the indirect cell-cell interactions among skin cells (keratinocytes) and immune cells (Dendritic, T Cells, and Macrophages) (Figure 1). Interactions depicted in the diagram elucidate the involvement of the IL-36 cytokine in the proliferation, maturation, differentiation, and apoptosis of modeled cells in a Psoriatic lesion. In a recent study, Pandey et al., (2021) have developed a mathematical model and demonstrated the involvement of cytokines TNF*α*, IL-23/IL-17, and IL-15 in the pathogenesis of psoriasis [4]. The mathematical modeling framework proposed by them would be ideal for the present study and therefore we follow that for modeling the network mediated by IL-36 which is as follows.

**Figure 1:**
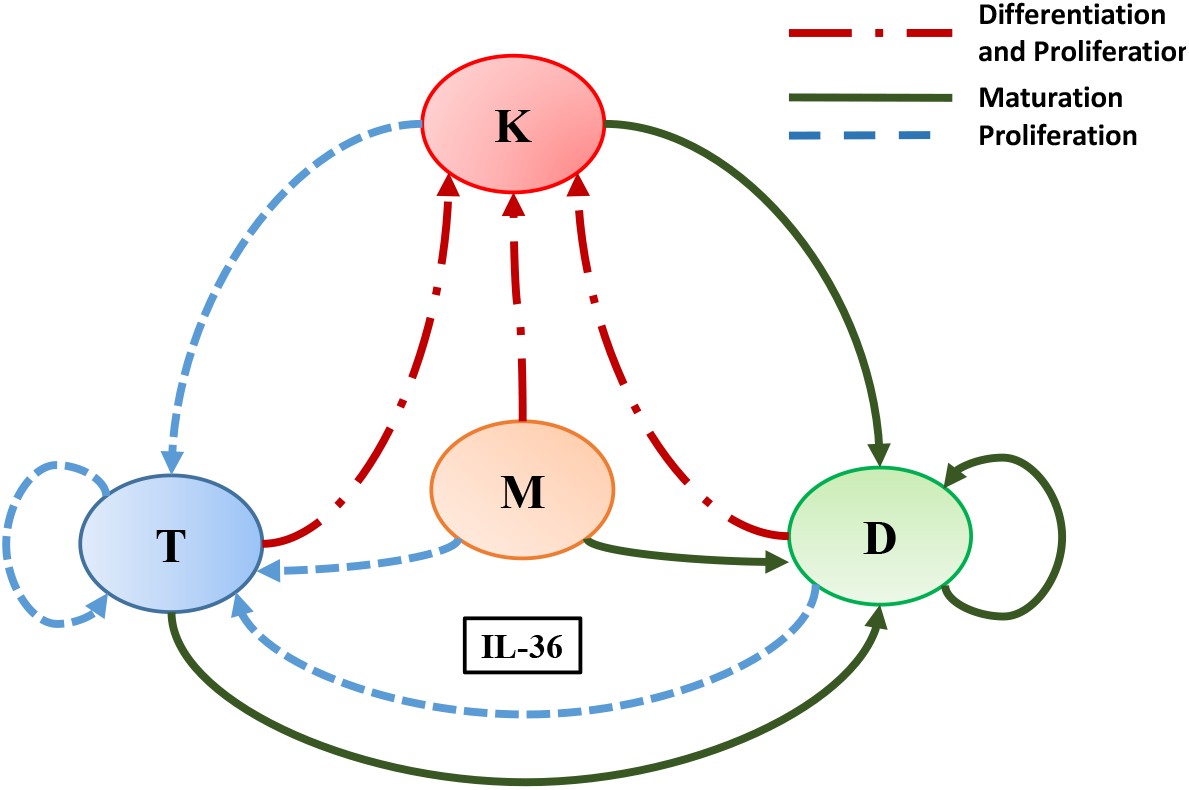
A schematic diagram explaining IL-36 mediated interactions among immune cells and keratinocytes involved in the pathogenesis of Psoriasis. Here L, K, D, and M denote T-Lymphocytes, Keratinocytes, Dendritic cells, and Macrophages, respectively. Different coloured arrows represent different types of interactions amongst the cells involved.

### 2.1 Mathematical Modelling Framework

In our model we consider a lesional skin of a psoriatic patient with the resident population of keratinocytes and immune cells (T Cells, Dendritic Cells, and Macrophages). We assume that all the modelled cells can infiltrate in this lesion. This region of skin is considered to be treated with the different level of IL-36 in order to investigate the role of that in the plaque formation and progression of Psoriasis.

The dynamics of the modeled cell population is governed by following ordinary differential equations (ODEs), if the framework proposed elsewhere [4] is applied

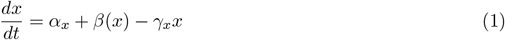

here *x* represents the population of a modeled cell type i.e. *x* = {*K, D, T, M*} where the population of keratinocytes, dendritic cells, T cells, and macrophages cells are denoted by K, D, T, and M, respectively. The parameter *α*_*x*_ denotes a fixed rate of increase in the population of modeled cells through their migration towards the lesion or differentiation/ maturation of them in the lesion (independent of the cytokine IL-36). The net rate of change in the cell population represented by the function *β* (*x*) which accounts for the cumulative effects of the cytokine in cell differentiation, maturation, proliferation, and apoptosis. The rate of decrease in the cell population through other processes including apoptosis not mediated by IL-6 is denoted by the parameter *γ*_*x*_. The function *β* is considered to be a function that saturates at *β*_*x*_ and is dependent on the population of all modeled cells or a few based on the interaction network shown in Figure 1. This dependence is modeled through the following two forms of *β*.

i. when *β* increases due to the modulatory effect of the population of all cell types or a few

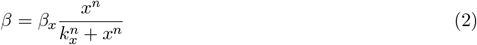
ii. when *β* decreases because of the modulatory effect of the population of all cell types or a few

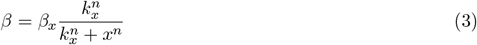

In Eqs.(2) and (3), *β*_*x*_ represents the saturated rate of change in the parameter *β* due to the modulatory effect of modeled cells. Here, the parameter *k*_*x*_ represents the population of a modeled cell type that is required to show the half maximal change in the population of either its own or in others depending on the interaction of the network shown in Figure 1. The slope of the function *β* is determined by the Hill coefficient-like parameter *n*. The rationale behind using Hill-like function here is based on the wide acceptance of that as an empirical saturating function.

### 2.2 Indirect Cell-Cell Interaction Network mediated by the Cytokine IL-36

All modeled indirect cell-cell interactions driven by IL-36 have been depicted in Figure 1 in the form of a schematic network. The following system of ordinary differential equations will be obtained if the above-mentioned modeling framework is applied to model this interaction network

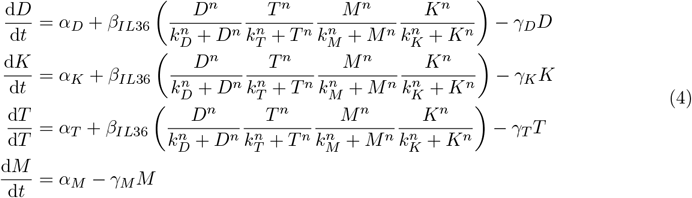

where D, K, T, and M denote the population of dendritic cells, keratinocytes, T cells, and macrophages cells, respectively. The parameter *β*_*IL*36_ represents the level of cytokine IL-36. The parameter *k*_*D*_ represents the population of dendritic cells in the lesion required to attain half the maximal effect of dendritic cell (D) on D, K, and T (illustrated in the network (1)). Similarly, *k*_*K*_, *k*_*T*_, and *k*_*M*_ denote the population of keratinocytes, T cells, and macrophages that are required for half the maximal effect of the keratinocytes (K), T cell (T), and macrophage (M) populations on dendritic cells, keratinocytes, and T cells, respectively. The parameters *α*_*D*_, *α*_*K*_, *α*_*T*_ and *α*_*M*_ represent the rates of migration (in cells/day) of dendritic cells, keratinocytes, T cells and macrophages, respectively to the site of psoriatic lesion. Their rates of apoptosis (in *day*^−1^) is represented by *γ*_*D*_, *γ*_*K*_, *γ*_*T*_, and *γ*_*M*_,respectively. The aforementioned set of ODEs indicates that the dynamics of macrophages population (M) are independent of the population of other cell types (D, K, and T). Hence, a quasi-steady state is assumed for macrophages and the dynamics of the system of ordinary differential equations is mainly governed by the following system of equations.

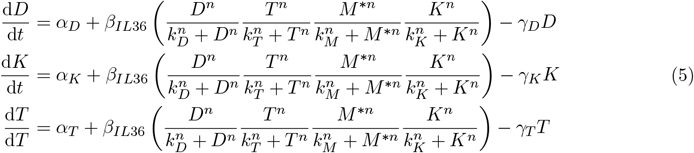

here, 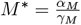 denotes the cell population of macrophage at steady-state. To further reduce the complexity of the system of ODEs, we make the system dimension less by assuming 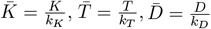, and 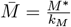. The dimension-less ODE system reads as

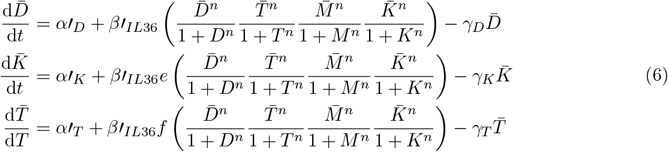

were, 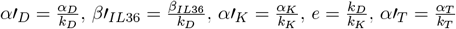 and 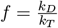.

### 2.3 Model Parameters

The typical values of all the model parameters used for obtaining numerical solutions of the system of ODEs are summarized in Table 1. The values for the rate of apoptosis of T cells (*γ*_*T*_), dendritic cells (*γ*_*D*_), and keratinocytes (*γ*_*K*_) are taken from a recent study [4].

**Table 1:** Default values of the model parameters.

| Parameters | Values | Biological Meaning |
| --- | --- | --- |
| $\alpha'_K$ | 0.1 day<sup>-1</sup> | Effective rate of migration of Keratinocytes towards the lesion |
| $\alpha'_D$ | 1 day<sup>-1</sup> | Effective rate of migration of Dendritic Cells towards the lesion |
| $\alpha'_T$ | 0.5 day<sup>-1</sup> | Effective rate of migration of T Cells towards the lesion |
| $\beta'_{IL36}$ | 4 day<sup>-1</sup> | Denotes the level of IL-36 Cytokine |
| $\gamma_K$ | 0.5 day<sup>-1</sup> | Rate of Apoptosis of Keratinocytes |
| $\gamma_D$ | 0.2 day<sup>-1</sup> | Rate of Apoptosis of Dendritic Cells |
| $\gamma_T$ | 0.5 day<sup>-1</sup> | Rate of Apoptosis of T Cells |
| $\bar{M}$ | 1 | Steady-state population of Macrophages |
| $n$ | 1 | Hill-like Coefficient |
| $e$ | 2 | Modulatory effect of Keratinocytes on other modeled cell types and itself |
| $f$ | 0.5 | Modulatory effect of Dendritic Cells on other modeled cell types and itself |

### 2.4 Bifurcation Analysis

To investigate the steady state behaviour of the model and how that changes for varying parameter values a Bifurcation analysis is performed using an open access tool MATCONT [18] and software MATLAB [19].

## 3 Results

We investigate the steady-state behavior of the network mediated by the IL-36 cytokine (Figure 1) using a standard bifurcation analysis technique of dynamical systems theory. IL-36 is considered to be a pro-inflammatory cytokine and its role in the progression of psoriasis has been observed [20]. Here, we study the effect of changes in the level of IL-36 cytokine on the population of keratinocytes for varying model parameters.

Our results suggest that an increase in the population of keratinocytes (K) and dendritic cells (D) would be observed for an increase in the level of the IL-36 cytokines (represented by 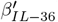) (Figure 2. A four-fold increase in the keratinocyte population is believed to be an indicator of a psoriatic condition based on the observation by Valeyev et. al [21]. Therefore, a significant increase in the level of IL-36 would lead to a psoriatic state which is in agreement with the observation that in psoriasis increase in IL-36 level has been observed [20]. In addition, a significant increase in the dendritic cell population has been observed in the pustule formation stage of GPP [22]. We observed that an increase in the value of the parameter 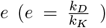 would cause a sharp increase in the keratinocyte population indicating a strong modulatory effect of keratinocytes on the other modeled cell types is strong compared to that of Dendritic Cells. Whereas, a gradual increase in the dendritic cell population is observed for an increase in the values of parameter *e* (Fig. 2A-B) which saturates for large values of *e*.

**Figure 2:**
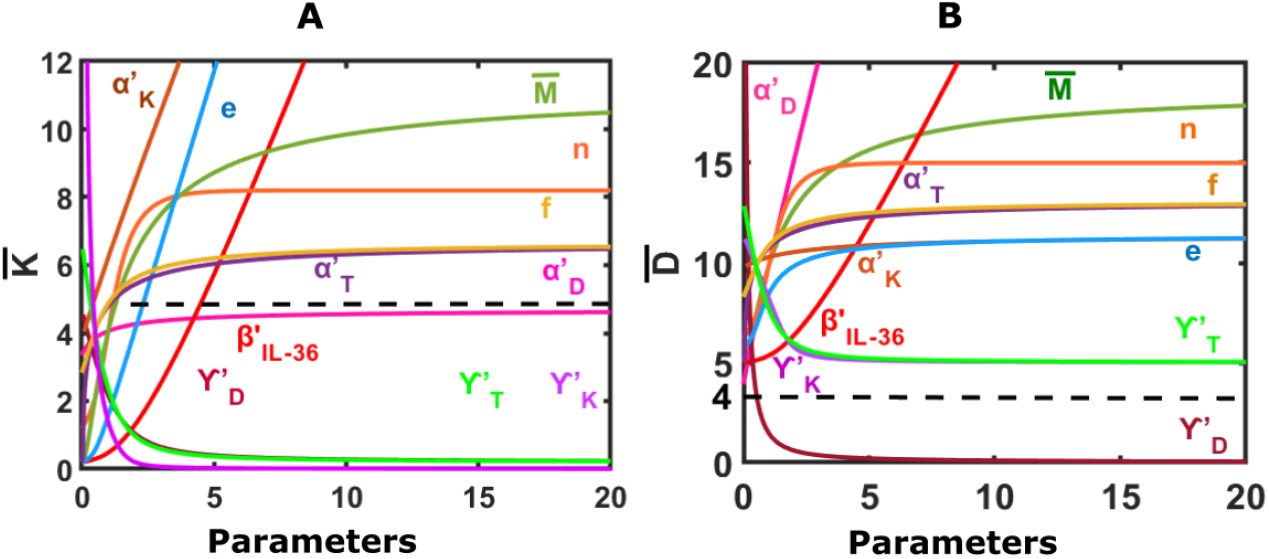
Steady state response of model for changes in different model parameters. (A) Change in the population of Keratinocytes for varying model parameters. (B) Change in the population of Dendritic Cells for varying model parameters. Default parameters are: 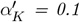, β_IL−36_ = 4, n = 1, 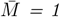, γ_K_ = 0.5, γ_D_= 0.2, γ_T_ = 0.5, 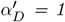, e = 2, 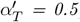, f = 0.5. Initial condition is: K = 4, D = 2, T = 0.1

Also, the keratinocyte and dendritic cell population would increase in the psoriatic lesion with an increase in the effective rate of migration of keratinocytes, dendritic cells, T-cells, and macrophages (represented by 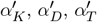, and 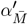, respectively) as shown in Figure 2A.

Further, an increase in the values of parameter 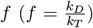 would result in a gradual increase in the population of keratinocytes and dendritic cells which saturates for further increase in *f* (Fig.2A-B). This suggests that the modulatory effect of the dendritic cell population on other modeled cells and itself is weak compared to that of T-cells.

In addition, our results suggest an increase in the Hill-coefficient-like parameter *n* would lead to an increase in the keratinocyte and dendritic cell populations initially that saturates at large values of *n*.

For an increase in the steady-state population of macrophages 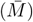, the population of keratinocyte and dendritic cells both increases gradually (Figure 2).

Through Figure 3, we demonstrate that the change in the population of keratinocytes upon varying the level of IL-36 (represented by *β*_*IL*−36_) depends on other model parameters. Our model results suggest a sharp increase in the keratinocyte population for an increase in the effective rate of inward migration of keratinocytes 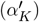 as depicted in 3A. A switch-like increase would be observed for small values of 3A. The bistable region bounded by the limit points would decrease with the increase in the 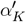. A sharp increase in the keratinocytes population would also be observed for an increase in the effective rate of inward migration of dendritic cells 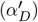 and the effective rate of inward migration of T -cells 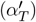 (Figure 3B and SF1). Also, in the case of a switch-like increase in the population of keratinocytes, the bistable region would decrease with the increase in 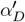 and 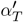 (Figure 3B and SF1).

**Figure 3:**
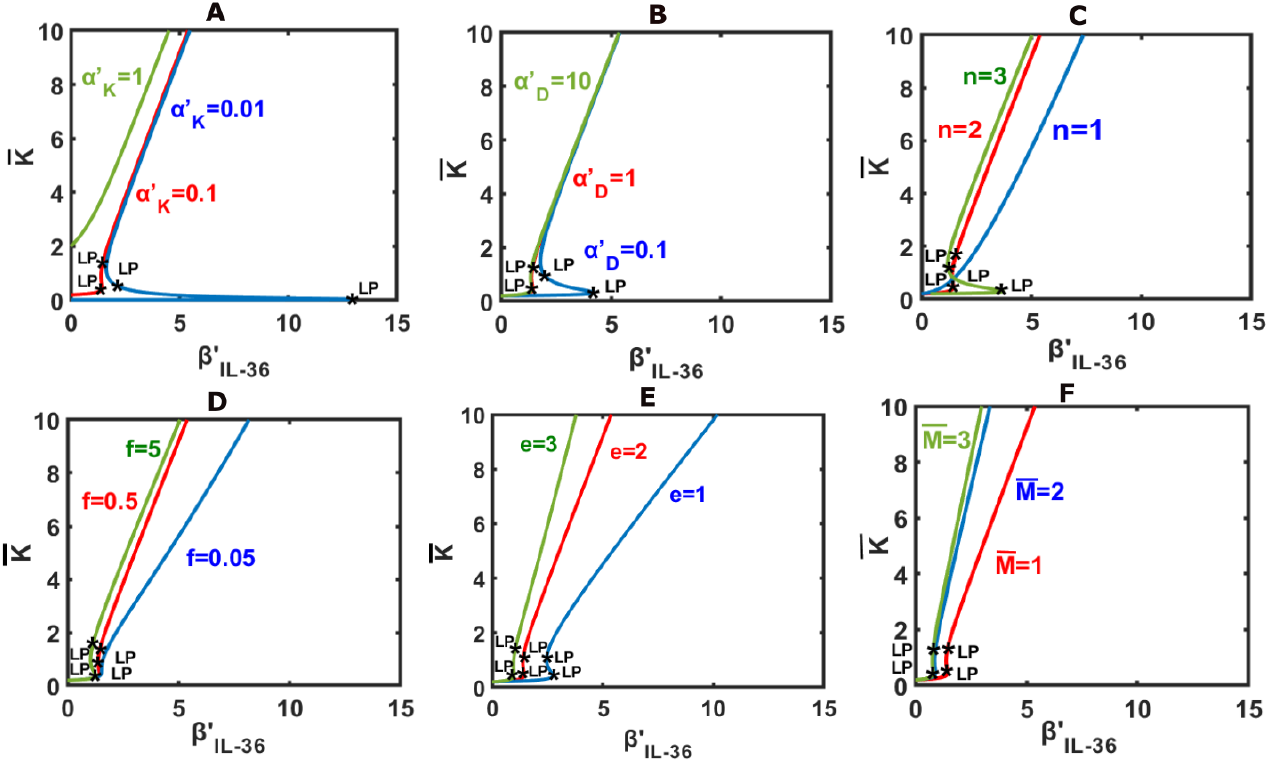
Keratinocyte population as a function of the level of IL-36 denoted by β’_IL−36_ (A) Change in the population of varying level of IL-36 for different values of the parameter 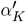. (B) Change in the population of varying level of IL-36 for different values of the parameter 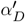. (C) Change in keratinocyte population as a function of the level of IL-36 for different values of n. (D) Change in keratinocyte population as a function of the level of IL-36 for different values of f . (E) Change in keratinocyte population as a function of the level of IL-36 for different values of e. (F) Change in keratinocyte population as a function of the level of IL-36 for different values of 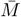. Default parameter values are: 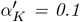, β_IL−36_ = 4, n = 2, 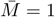, γ_K_ = 0.5, γ_D_= 0.2, γ_T_ = 0.5, 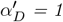, e = 2, 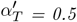, f = 0.5. Initial condition is: 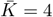, D = 2, T = 0.1

Further, a decrease in the population of keratinocyte population would be observed for an increase in Hill-like-coefficient (Figure 3C). For a small value of *n*, the increase in the population of keratinocytes would occur gradually. A switch-like increase is expected to occur for a large value of *n* and the bistable region would be larger for *n* = 3 in comparison with *n* = 2.

Our results suggest a gradual and switch-like increase in the population of keratinocytes for an increase in values of *f* (denoting the modulatory effect of keratinocytes on itself) and *e* (denoting the modulatory effect of dendritic cells on other modeled cell types (Figure 3 D and E). Similar observations would be observed for an increase in the steady-state population of macrophages 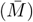 (Figure 3F).

The switch-like increase in keratinocytes would be a robust behaviour as we found large bistable regions in two-parameter space spanned by (*n*, 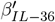) (Figure 4 A). A typical temporal behaviour of the network in bistable and monostable region is shown in Figure 4 B and C.

**Figure 4:**
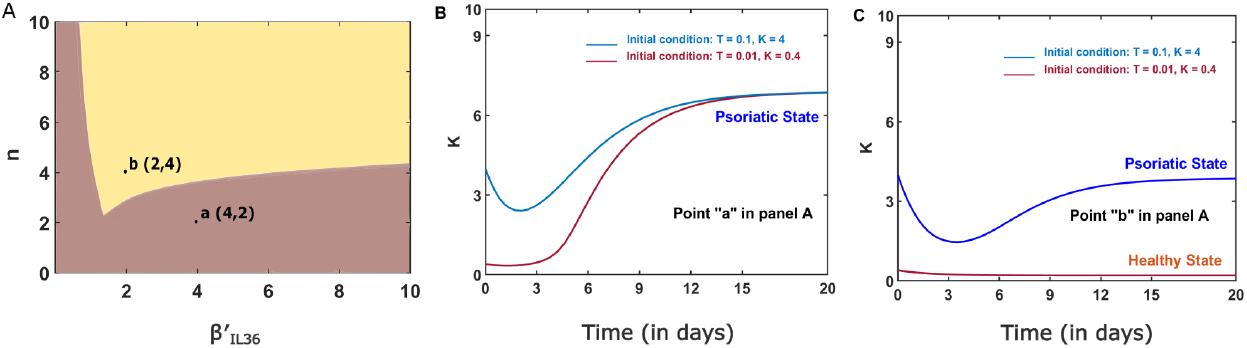
(A) Two Parameter Bifurcation Curve for n and 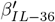.(B) Dynamics of the network for a model parameter combination that lies inside a bistable region in the panel Figure 3A.(C) Dynamics of the network for a model parameter combination that lies inside a monotstable region in the panel. Parameters and initial condition is same as in Figure3

## 4 Discussion

Psoriasis is characterized by the formation of a psoriatic lesion that is infiltrated by immune cell populations [2, 14]. Several mathematical models have been developed to elucidate the role of cytokines in the progression of Psoriasis [21, 23, 24]. In one of the model the role of IL-15, IL-17/IL-23 and TNF*α* cytokines mediating indirect cell-cell interactions amongst Keratinocytes, Dendritic Cells and T-Cells in a psoriatic lesion has been explored [4]. Obvious targets for IL-15, IL-17/IL-23 and TNF*α* cytokines have been translated into drugs that inhibit them.

Different types of Psoriasis have been reported and Psoriasis vulgaris, are considered as the most prevalent type amongst the patients. General Pustular Psoriasis (GPP) is another severe form of Psoriasis. In GPP, relapsing sterile pustules are present in the affected individuals [9]. Recent studies attribute this relapse to T cells (skin resident memory *T*_*RM*_ cell, memory like *γδ*T cells) whose proliferation and differentiation is caused by dendritic cells through a feed-forward loop involving inflammatory cytokines (reviewed in [25]). The dermal dendritic cells colocalizes with neutrophils secreting elastase which is believed to have a role in pustule formation [26]. Studies have shown that the presence of Human Neutrophil Elastase is elevated in GPP that contributes to tissue damage, degradation of extracellular matrix and disruption of skin barrier function [5]. Hence, the increase in the level of DCs and its indirect interaction with elastase [27] could lead to the worsening of psoriasis. Recent studies have unveiled IL-36 cytokine as a potential drug target for GPP/Psoriasis [28–30].

This hypothesis is augmented by a gene expression data that reports a significant upregulation of IL-36 cluster of cytokines in Psoriasis patients [**?**]. To explore the role of pro-inflammatory cytokine IL-36 in the plaque formation and disease progression of Psoriasis, we have built a network of indirect cell-cell interactions among key immune cells (Dendritic cells, Macrophage, and T-lymphocytes) and skin cells (Keratinocytes) mediated by IL-36. Our results are consistent with the finding that IL-36 could lead to worsening of diseased state in Psoriasis. Our results suggest an increase in the population of keratinocytes; thus the plaque formation and progression of Psoriasis could occur either in a gradual or switch-like manner. The switch-like-on set or progression of Psoriasis would occur due to the bistable behaviour of the network formed due to the indirect immune and keratinocytes cell interactions via IL-36. Parameters controlling the bistable region have been explored that suggests the switch-like abrupt changes in the keratinocyte population is a robust phenomenon. For instance, we found a sizeable bistable region in two-parameter space spanned by the level of IL-36 cytokines 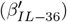 vs Hill-coefficient like parameter (*n*).

One of the limitations of our model is that it is formulated for only a single cytokine whereas the role of several cytokines in Psoriasis is well-reported. Here, it is assumed that although there are several cytokines involved in Psoriasis, IL-36 has a dominant role, especially in GPP. In addition, model results are yet to be experimentally verified. However, the present framework could be used to study the role of a dominant cytokine in other inflammatory and autoimmune diseases.

## Author Contributions

RP and RKM conceived the idea. SP and ST performed simulations. RP, SB, ST, and SP analyzed the simulation results. All authors contributed to the preparation of the manuscript. All authors have read and agreed to the published version of the manuscript.

## Acknowledgments

RP acknowledges the Department of Science and Technology, India for the DST-INSPIRE Faculty Award (DST/INSPIRE/04/2015/001939) and Banaras Hindu University for the IoE seed grant. RP also acknowledges the University Grant Commission, India start-up grant awarded to him. SB acknowledges the Prime Minister Research Fellowship awarded to her by the Ministry of Education Government of India. ST acknowledges Banaras Hindu University for the doctoral fellowship.

## Conflicts of Interest

The authors declare that the research was conducted in the absence of any commercial or financial relationships that could be construed as a potential conflict of interest.

## Notes

### Competing Interest Statement

The authors have declared no competing interest.

### Summary of Updates

To add equal contribution notation by following authors in the HTML version: Sneha Pandey*, Syona Tiwari* and Sulagnna Basu*. No change has been made in the manuscript file.

