## Supplementary for "Modeling of the Network Mediated by IL-36 Involved in Psoriasis"

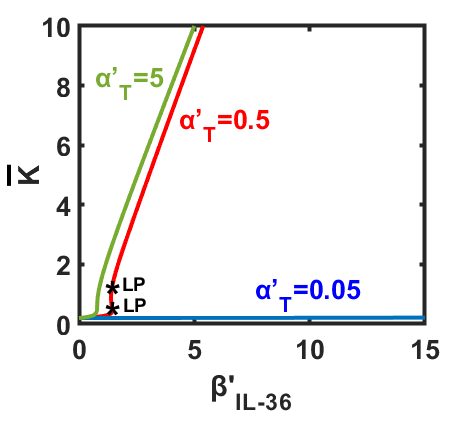

**Figure 1:** Keratinocytes vs. ${\beta'}_{IL36}$: Comparative graph for varying levels of Effective rates of migration of T- cells towards psoriatic lesion.
